# Integrative multi-omic analysis identifies key transcription factors and target proteins in renal cell carcinoma and its subtypes

**DOI:** 10.1101/2025.11.23.690024

**Authors:** Surya B. Chhetri, Timothy D. Winter, Mitchell J. Machiela, Kevin Brown, Alexis Battle, Mark P. Purdue, Stephen Chanock, Diptavo Dutta

## Abstract

To characterize key transcription factors (TFs) whose differential DNA binding can be altered by genetic variants associated with risk for renal cell carcinoma (RCC), we conducted a series of mixed model-based analyses integrating 449 TF ChIP-seq profiles across 9 kidney-related cell lines and summary statistics from a multi-ancestry genome-wide association study of RCC. We identified 96 unique TFs for which presence of SNPs in a neighborhood of TF ChIP-seq peaks are significantly associated (p-value <1x10-4) with their effect on RCC, including EPAS1, ARNT, PAX8 and PBRM1, previously implicated in RCC pathogenesis. Most TFs overlapped active promoters/enhancers in RCC tumors but remained significant after adjusting for tumor chromatin accessibility. Further, we found the co-occupancy of 220 pairs of RCC-related TFs to be associated with RCC risk (FDR<5%) beyond effects of individual TFs, highlighting synergistic regulation between pairs of TFs. To further investigate distal (trans) regulation of TF-binding disruption at RCC associated loci on the proteome, we used a set-based regression to aggregate the trans-effects of multiple loci overlapping with TF binding sites. Across 2,732 proteins profiled in UKB-PPP, identified 169 trans-associated (p-value<1.6x10-7) proteins, nominating specific targets for each TF. For example, we identified TLR3 and ZP3 to be associated with EPAS1, ARNT, and PBRM1, indicating these proteins are likely affected by RCC-related variants disrupting binding sites of the corresponding TFs. These results characterize the landscape of RCC-related TFs and implicate TF-mediated proteomic mechanisms in RCC pathogenesis, nominating testable targets for laboratory studies.

## Introduction

Kidney cancer represents a significant global health challenge with increasing incidence and mortality rates, especially in Western populations^1^. Over 85% of kidney cancers are renal cell carcinoma (RCC), which includes several histologic subtypes, most commonly clear-cell (ccRCC; over 75% of RCC cases), papillary (papRCC; 15%), and chromophobe (chrRCC; 5%) carcinomas^2^. In addition to several factors, such as obesity^3^, smoking^4^, and hypertension^5^, recent large-scale genotyping efforts have revealed a substantial role of inherited common genetic variation in RCC etiology based on recent genome-wide association studies (GWAS)^6,7^ that identified have 108 risk variants across 63 susceptibility regions. However, the majority of RCC GWAS loci reside in non-coding regulatory regions and are postulated to influence RCC risk through dysregulation of gene expression that leads to perturbations of specific cancer pathways, as shown in detailed analysis of loci that are involved in apoptosis, cell growth or SWI/SNF activity^8–10^.

Differential alternation of transcriptional regulation by sequence-specific DNA-binding factors, chromatin modifiers, and transcriptional co-regulators which we collectively refer to as transcription factors (TFs), is a common feature of risk loci identified by GWAS for mapping complex diseases^11–13^ such as cancer. Investigation of the regulatory landscape surrounding GWAS loci can identify dysregulated gene expression profiles that influence tumor initiation, progression, and metastasis^14,15^. Post-GWAS *in silico* analysis, followed up with functional experiments, have suggested that regulatory risk variants can disrupt DNA binding affinities of TFs, and thus play a key role in mediating cancer risk^16^. By binding at specific DNA motifs and interacting with co-activators or co-repressors, TFs can regulate the transcription of genes that control key cellular processes, such as proliferation, apoptosis, and differentiation^17^. In complex diseases like cancers, TFs in conjunction with genetic variants at risk loci can regulate downstream biological networks, offering new ways to understand how genetic susceptibility translates into cancer biology. Furthermore, the combinatorial action of TFs and their co-occupancy at specific genomic loci can further amplify or modulate effects, highlighting the complexity of their roles in transcriptional regulation. Studying the impact of TF binding on gene expression and its effect on cancer risk can contribute to identifying novel therapeutic targets and understanding the biology of cancer susceptibility. For example, studies have shown pathogenic dysregulations of gene expression are mediated through risk variants altering binding affinities of master TFs, such as *FOXA1*, *ESR1*, and *AR* for breast and prostate cancers^14,18,19^.

Several TFs have been implicated in RCC pathogenesis through functional studies of candidate TFs, including HIFs (hypoxia-inducible factors), *PAX8, FOS*, and *JUN*^20–24^. HIFs, which are stabilized in the hypoxic tumor environment of ccRCC due to mutations in the *VHL* gene, regulate a network of genes involved in angiogenesis, metabolism, and proliferation^25–27^. *PAX8*, a lineage-specific TF essential for kidney development, has been identified as a key oncogenic driver in RCC^23,24^. *FOS* and *JUN*, components of the AP-1 complex, are involved in stress responses and have been shown to influence RCC progression^28,29^. Analyses in specific GWAS identified loci such 8q24.21^8^, 11q13.3^30^, 12p12.1^9^, and 14q24^10^, have identified important contributions of HIFs, and SWI/SNF complexes and identified target genes in these loci for RCC susceptibility. Despite these discoveries, an agnostic evaluation of the genome-wide contribution of these and other TFs on RCC remains unexplored. Moreover, the role of TF co-occupancy and combinatorial regulation, which could refine the transcriptional programs driving RCC susceptibility, is yet to be investigated as well.

In RCC, as in other cancers, disruptions in TF binding often can affect key cellular proteins involved in proliferation, apoptosis, metabolism, and immune modulation^13,31^. A recently proposed “omnigenic” model postulates that effects of genetic variants associated with complex diseases, including cancer, are mediated via an intricate intermediate network resulting in *trans-*QTL effects on a few “core genes”^32,33^. If such an “omnigenic” characterization is reflective of the genetic architecture of RCC, then identifying the potential genes and proteins *trans*-associated with TF binding disruptions at RCC-associated GWAS loci can help prioritize putative disease-relevant molecular targets. However, a major challenge has been the limited statistical power for detection of *trans-*associations^34,35^ due to possibly weaker effect sizes as well as the lack of innovative approaches to aggregate such weaker effects of multiple variants converging on similar molecular targets. As a consequence, systematic studies connecting GWAS risk loci, TF activity, and downstream protein-level consequences in RCC remain limited, despite their critical role in elucidating mechanisms relevant to RCC.

In this study, we conducted a genomic and epigenomic analysis to identify the TFs and potential target proteins driving RCC susceptibility using the largest GWAS of RCC and its subtypes to date^7^, coupled with publicly available chromatin immunoprecipitation followed by high throughput sequencing (ChIP-seq) data from 449 TF ChIP-seq experiments across diverse kidney-related cell types^36,37^. We refined the association of TF-occupancy patterns with risk of RCC and its two major histologic subtypes, ccRCC and papRCC by integrating histone modification profiles of RCC tumors. Additional analyses were conducted to identify co-occupancy patterns of TFs that were associated with disease risk beyond the effect of individual TFs. Finally, using large-scale proteomic profiling in UK Biobank^38^, we identify potential downstream target proteins which might be dysregulated by the disruptions in TF binding at RCC-associated loci.

## Results

### Overview of Methods

#### (A) Association of TFs with RCC and its subtypes

To study how genetic variations at TF binding sites impact RCC risk, we applied a linear mixed model integrating TF ChIP-seq data with GWAS summary statistics, while controlling for local linkage disequilibrium (LD) as random effects. LD blocks were defined using non-overlapping 1Mb segments and were considered to be independent (See **Methods**). Subsequently, we adopt a previously used mixed effect model^39^ to identify the association between a given TF and risk of RCC (or its subtypes) as:

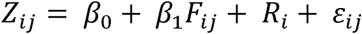

where Z_*ij*_ is the probit transformed RCC (or its subtypes)-association p-value for the j^th^ SNP in the i^th^LD block, F_*ij*_ is the “occupancy” of the TF of interest, a variable taking values 1 when the j^th^ SNP in the i^th^ LD block overlaps with a ChIP-seq peak for the TF of interest and 0 otherwise; R is a random effect term for the i^th^ LD block, capturing within-block correlation. *β*_1_ is the effect of the given TF on RCC risk, which if significantly greater than 0 implies that SNPs in the ChIP-seq peaks for the TF of interest have larger z-scores and hence higher association with RCC, than expected by chance alone. This model is fitted for each TF separately to evaluate its association with RCC risk. Further, we repeated this analysis controlling for RCC tumor chromatin accessibility (CA; See **Methods**), to identify the TFs whose association with RCC risk was not solely due to genetic variants in CA regions. Additionally, we tested the effect of the co-occupancy of a pair of TFs, by incorporating a linear interaction term, representing the variants overlapping with ChIP-seq peaks of two TFs simultaneously (See **Methods**).

### (B) trans-effects of TF occupancy at RCC-associated loci on the proteome

To investigate the downstream effects of TF binding disruption at RCC-associated GWAS loci, we integrated plasma proteomic data from the UK Biobank Pharma Proteomics Project (UKB-PPP)^38^, encompassing 2,940 proteins from 34,557 individuals of European ancestry. For each TF, we defined a "TF-specific subset" of variants by identifying RCC-associated variants, or their LD-proxies, overlapping with TF ChIP-seq peaks and multiplying them by corresponding GWAS effect sizes, resulting in a “TF-partitioned score” F_*sub*_. These scores represent the cumulative effect of TF binding disruption at “TF-specific subset” of variants on RCC risk. We model the association between these disruption scores and k^th^ normalized plasma protein level P, using a set-based regression approach:

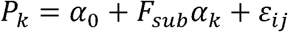

where α_*k*_ is a random effect of the TF-partitioned scores F_*sub*_. The test for α_*k*_ = 0 can be performed using a variance component score test^40–42^ under different kernel assumptions using proteomic summary statistics only (See **Methods**). A significant α_*k*_ indicates an association between the TF-partitioned scores for the TF of interest and the k^th^ protein level. In particular, we focus on the associations between a TF-partitioned score for a TF of interest, and all the proteins with a distal (> +/- 5Mb) transcription start site for each “TF-specific subset” of variants, reflecting the *trans*-effects of the SNPs. Thus, the *trans*-association between a TF-partitioned score and a distal protein intuitively represents the convergent *trans*-effect of TF binding disruption at a subset of RCC associated loci on the protein and hence can highlight potential downstream target proteins.

### Transcription factors associated with RCC and its subtypes

Across public resources like ENCODE, GEO and ChIP-Atlas, we curated a list of 449 TF ChIP-seq experiments performed in normal (N = 401) and tumor (N = 48) kidney cell lines, with number of peaks ranging from 114 to 382,880 across the genome (**Supplementary Table 1**). Using the linear mixed model approach described above (See **Methods**), we integrated these TF ChIP-seq data with the GWAS summary statistics from Purdue et al^7^ on RCC (N_cases_ = 29,020) to identify TFs significantly associated with the risk of RCC or its subtypes. At a p-value threshold of 1×10^−4^ (reflecting the Bonferroni correction threshold for 449 TF ChIP-seq experiments), we identified 103 associations with RCC risk across 96 unique TFs (**Figure 1A, Supplementary Table 1**). Of these, 19 were not identified by S-LDSC while 23 were not identified by bootstrap enrichment^21,43^ (See **Methods** and **Supplementary Figure 1**). The strongest associations were seen for *ZNF335* (p-value = 3.2×10^−18^) in HEK293, *CREB1* (p-value = 2.0×10^−16^) in kidney cortex and *SMAD4* (p-value = 2.5×10^−16^) in the 786-O cell line respectively, with several additional zinc-finger TFs exhibiting strong associations. Several TFs known to be associated with RCC or regulators of pathways central to renal carcinogenesis were also identified among the 103 RCC-associated TF ChIP-seq profiles. For example, we identify associations with *ARNT* (minimum p-value = 3.1×10^−9^, 1 out of 5 cell lines) and *EPAS1* (minimum p-value = 3.8×10^−9^, 4 of 5 cell lines) which are key components of the hypoxia-inducible factor (HIF) complex and *PAX8* (P = 8.2×10^−8^, 1 of 1 cell line), a lineage-specific TF essential for kidney development. Additionally, we identify *PBRM1* (P = 2.3×10^−7^, 1 of 1 cell line), a subunit of SWI/SNF chromatin remodeling complex that is frequently mutated in RCC and acts as a tumor suppressor^44–46^. Other associated TFs include *SMAD4* (P = 1.2×10^−8^), *EP300* (minimum p-value = 5.4×10^−7^), *TAF1* (P = 1.3×10^−6^), and *KAT6A* (P = 2.8×10^−6^), all of which have been implicated in processes related to multiple cancers^47–51^ and might represent candidate master regulators that integrate upstream risk signals to drive aberrant tumorigenic processes and in particular, RCC development. Overall, the 96 TFs associated with RCC were enriched in several protein complexes including MLL3/4 (P = 4.3×10^−5^), SWI/SNF (P = 4.7×10^−5^), ATPase (P = 2.3×10^−7^), NCOR1 (P = 2.3×10^−7^) and PBAF (P = 2.3×10^−7^) among others, across different databases (**Supplementary Table 2**).

**Figure 1:**
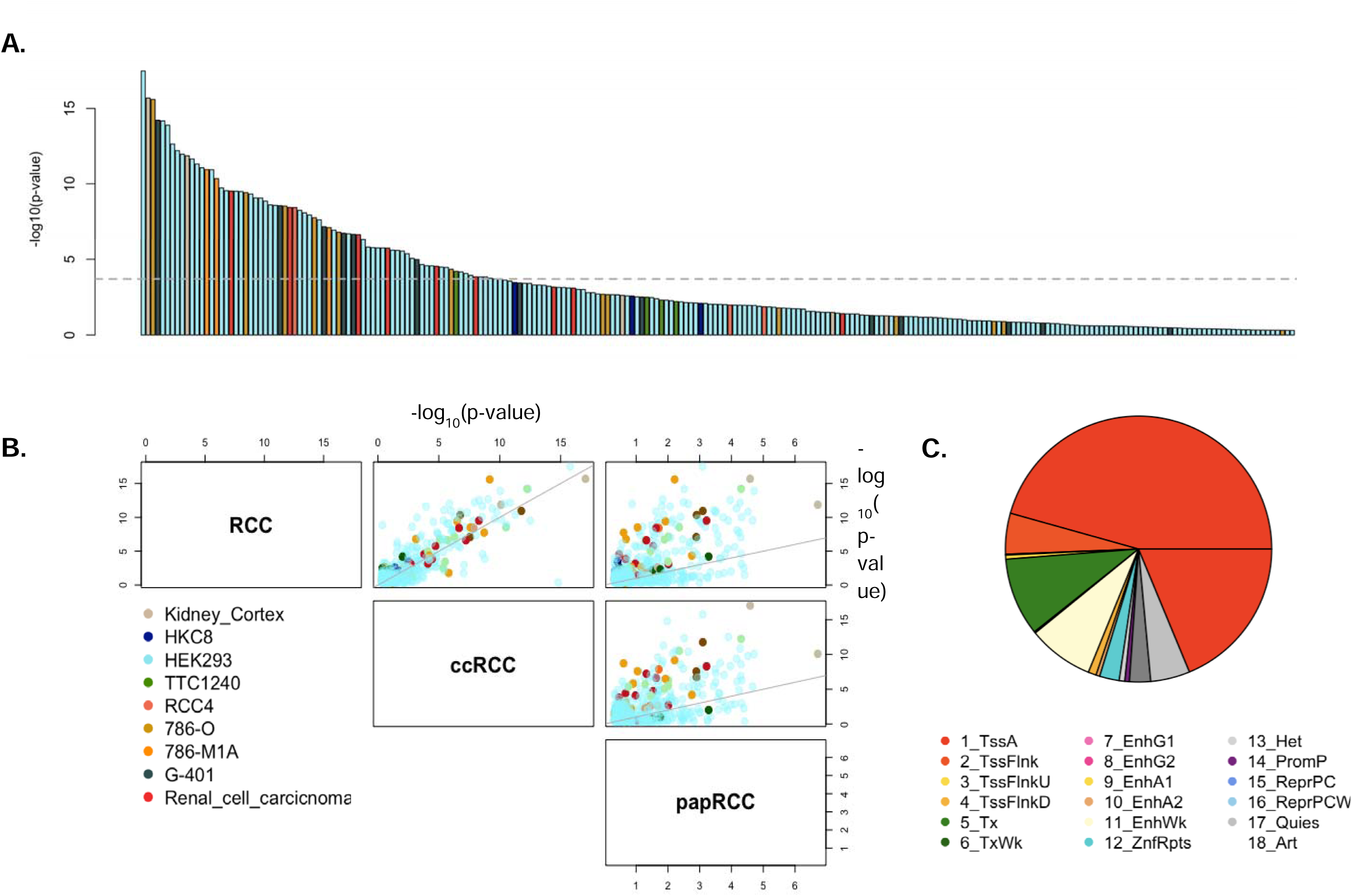
Association of TFs with RCC and subtypes. (A) Barplot showing -log10(p-value) of association of all 449 TF ChIP-seq profiles with RCC, colored by the respective cell line. The dashed line marks the p-value significance threshold of 1.0×10^−4^. (B) Comparison of associations (-log10 p-value) across all TF ChIP-seq profiles, for RCC and its subtypes (ccRCC and papRCC) colored by the respective cell line. (C) Epigenomic annotation of TF ChIP-seq peaks in kidney cell from ROADMAP using ChromHMM 18-state model depicting the proportion of peaks, averaged across all TFs, in each annotation.

Our findings suggest that common genetic variants at the binding sites of the RCC associated TFs cumulatively may potentiate RCC susceptibility. To further support potential TF binding alteration, we pursued a limited exploration of allele-specific binding patterns at RCC-related GWAS significant loci among TFs in HEK293 with significant occupancy and co-occupancy effects. We identified 7 TFs with nominally significant allele-specific binding effects, including four with allele-specific binding at more than one locus (See **Methods**; **Supplementary Table 3**). For example, PRDM6 (P = 8.7×10^−6^) ChIP-seq profile in HEK293 exhibited a highly significant (P = 2.91×10⁻¹¹) preference for “C” allele of rs4869977 (6q25.1) over “G” allele, the latter being the risk allele, indicating variation from C to G at this location is likely to disrupt *PRDM6* binding. We identified two additional associations with PRDM6 binding at different loci (rs11085218 at 19p13.2; rs141379009 at 11q22.3), along with two loci for ZNF121, ZBTB6 and KLF8 also displaying significant imbalance, thus suggesting potential allele-specific regulatory activity of TFs.

In analysis of the subtypes, ccRCC (N_cases_ = 16,321) and papRCC (N_cases_ = 2,193), we found 96 and 17 TF ChIP-seq profiles associated respectively, of which 19 and 3 were not identified for overall RCC. Association of TFs with ccRCC was largely concordant with that of RCC (**Figure 1B**) as expected given that the majority of cases in the RCC GWAS were of the ccRCC subtype. Of the 19 ccRCC TFs, not associated with RCC, we identify several known TFs that have been previously implicated in ccRCC tumorigenesis. For example, *BAP1* (P = 4.4×10^−5^), *JMJD6* (P = 1.6×10^−6^) and multiple members of Kruppel-like factor family like *KLF1* (P = 9.4×10^−5^) and *KLF9* (P = 1.7×10^−9^) were found to have a stronger association with ccRCC, although, as expected, their association with RCC was suggestive as well (P < 0.05). In particular, *JMJD6* which was assayed both in tumor as well as normal-like cell lines, exhibited association only in tumor cell line 786-O but not in normal embryonic cell line HEK293 (P = 0.04) indicating that the variants at its binding sites, specifically in tumors, has a significant aggregated effect on ccRCC. In contrast, papRCC exhibited a more distinctive pattern of association (**Figure 1B**) with the TFs although this analysis had lower statistical power given the comparatively limited number of papRCC cases in the corresponding GWAS (N_cases_ = 2,193). *RBAK* (P = 3.9×10^−5^), *SCRT2* (P = 4.8×10^−5^), and *ZBTB12* (P = 4.1×10^−5^) were associated to papRCC only, of which *RBAK* did not have suggestive associations with ccRCC or RCC (**Supplementary Table 1**).

### Adjusting for tumor chromatin accessibility

To evaluate the robustness of the above results, we further refined the association between TF ChIP-seq profile and RCC risk, by adjusting for tumor chromatin accessibility (CA). Using CA profile defined by ATAC-seq in TCGA-KIRC^52^, we found 76 TF ChIP-seq profiles (out of 103 previously identified) to be significant at 1×10^−4^ while 99 were significant at a nominal threshold of p-value < 0.05 (**Figure 2A**). Using CA profiles in RCC tumors defined by ATAC-seq in Nassar et al^53^, we found 76 previously identified TF ChIP-seq experiments to be significant at 1×10^−4^ while 92 were significant at a nominal threshold of p-value < 0.05 (**Supplementary Table 4**). We found that 61 unique TFs were consistently significantly associated (p-value < 1×10^−4^) with RCC risk across both analyses, including known TFs like EPAS1, PAX8 and ARNT. Thus, despite the overlap between CA and ChIP-seq peaks, majority of the TFs have effect on RCC beyond general CA in tumors.

**Figure 2:**
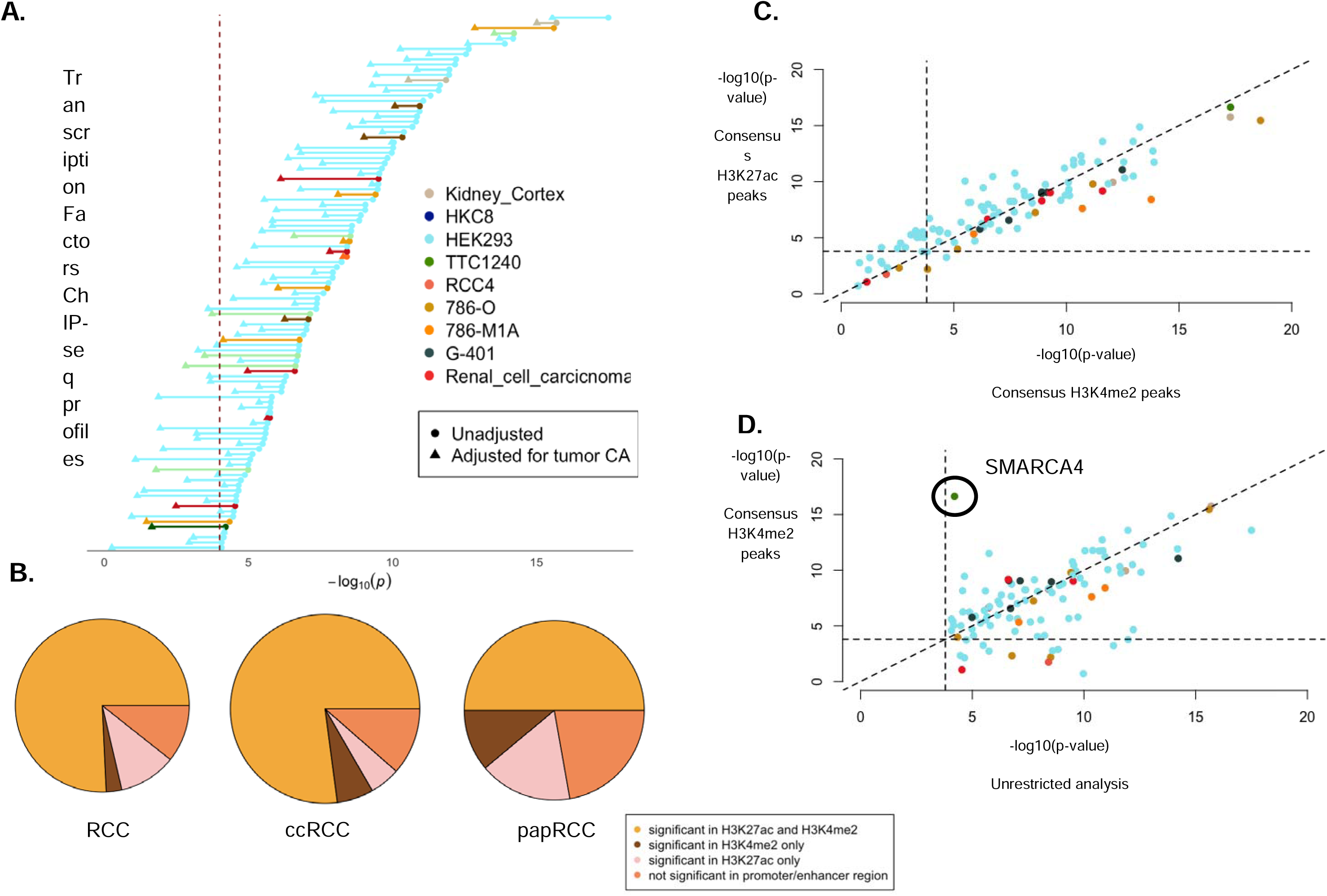
Integration with tumor epigenomic profiles. (A) Dumbell plot showing comparison between association of TF ChIP-seq profiles and RCC risk, using unadjusted model and adjusted for tumor CA reported by Nassar et al. (See **Results** and **Methods**) (B) Pie charts showing proportion of associated TFs that remained significant after restricting to active regions marked by H3K4me2 and H3K27ac profiles in RCC tumors through consensus peaks (See **Results** and **Methods**). (C) Comparison of association (-log10 p-value) of TFs with RCC, restricted to consensus peaks of H3K27ac (vertical axis) wi3th that of H3K4me2 (horizontal axis) colored by the respective cell lines. (D) Comparison of association (-log10 p-value) of TFs with RCC, restricted to consensus peaks of H3K4me2 (vertical axis) with unrestricted analysis (Figure 1A). SMARCA4, showing stronger association in the former is marked.

### Integration of risk TF binding with RCC tumor epigenomic profiles

Genomic annotation of the ChIP-seq profiles of the TFs identified to be associated with RCC or its subtypes revealed significant enrichment at regulatory elements marked by active chromatin states in normal fetal kidney epigenomes from ROADMAP^54^ (**Figure 1C**; See **Methods**), as we expected. To delineate the regulatory associations of these TFs, we repeated the above analysis by further restricting the “occupancy” of TF binding events to active regulatory regions in RCC tumors specifically. Leveraging comprehensive histone modification (H3K4me2 and H3K27ac) profiles across different stages, grades and subtypes of RCC tumors^53^, we first defined two distinct categories of regulatory regions: (1) “consensus peaks,” representing chromatin regions consistently marked by respective histone modification patterns, indicative of a shared regulatory landscape across RCC tumors, and (2) “union peaks,” encompassing both shared and unique tumor epigenomic patterns across tumors. Of the 103 RCC-TF associations, 83 (80.58%) remained significant within “consensus peaks” of H3K4me2, which marks regulatory elements poised for or engaged in transcriptional activity, while 91 (88.35%) remained significant when restricted to “consensus peaks” for H3K27ac, a marker of active regulatory elements, particularly enhancers (**Figure 2B**), with 77 (74.8%) associations significant in both (**Supplementary Table 5**). Association results were qualitatively concordant between consensus peaks and union peaks, the later representing a larger variability in regulatory potential. Curiously, TFs in developmental cell lines (HEK293) showed marginally stronger association at H3K27ac peaks (**Figure 2C**). We observed that compared to the previous analysis, associations of 52 (50.49%) and 45 (43.69%) TFs were stronger at H3K4me2-marked and H3K27ac-marked regions respectively, although we did not perform any systematic test of differences in effects. In particular, in the TTC1240 cell line *SMARCA4*, a core ATPase component of the SWI/SNF chromatin remodeling complex, showed notable stronger associations at both consensus H3K4me2 (P = 2.3×10^−17^) and H3K27ac peaks (P = 5.3×10^−18^) compared to previous analysis of only ChIP-seq peaks (P = 6.2×10^−05^), highlighting its regulatory potential in RCC (**Figure 2D**). In subtype-specific analysis, for ccRCC, we identified 82 (85.42%) TF ChIP-seq profiles exhibiting significant associations at both H3K4me2-marked and H3K27ac-marked consensus regions, while papRCC resulted more limited TF ChIP-seq associations, with only 7 (41.18%) and 8 (47.06%) TF ChIP-seq profiles showing significant associations at these respective regions.

### Co-occupancy of TFs reveal synergistic regulation

We investigated whether variants at binding sites of multiple TFs jointly conferred additional disease risk beyond the individual effect of a single TF binding events by analyzing genomic regions co-occupied by multiple TFs. Of the initially identified set of 103 RCC-risk-associated TF ChIP-seq profiles, our analysis focused on 77 risk-associated TF ChIP-seq profiles with significant associations in both H3K4me2-marked and H3K27ac-marked regulatory regions using an interaction model (See **Methods**). We identified 220 significant pairwise TF interactions at an FDR < 5% (10 are significant at Bonferroni threshold of 1.5×10^−5^) representing potentially synergistic regulation by multiple TFs, in addition to the effect of individual TFs (**Supplementary Table 6)**. The strongest associations were identified in HEK293 cell lines, to be between *ZNF324* and *ZNF121* (P = 1.2×10^−7^) and between *PRDM6* and *KLF8* (P = 2.4×10^−7^) indicating additional effects of the variants co-occupied by each of the TFs beyond their individual effects (**Figure 3**). 82% of the associations were found to be nominally significant at p-value < 0.05 after adjusting for tumor chromatin accessibility (**Supplementary Table 6**; See Methods). Several of the identified interactions reflected previously reported interactions between the TFs. For example, we identify significant interaction between *BRD7* and *MYC* (P = 1.7×10^−5^) in the G-401 cell line, the former being a part of the SWI/SNF chromatin remodeling complex and can suppress the activity of the latter which is a known oncogene. Among the remaining, several significant interactions involved Zinc finger genes, between which are known to interact with chromatin modifiers, modulating critical RCC related gene programs. From existing protein interaction databases (STRING^55^, CORUM^56^ and BioGrid^57^) we found at least 34 pairs with evidence of physical interaction (**Supplementary Table 6**). In subtype-stratified analysis, we identified 27 significant interactions (FDR < 5%) for ccRCC of which 7 had evidence in existing interaction databases (**Supplementary Table 6**) while none were identified for papRCC. In particular for ccRCC, we identified significant interactions of *PBRM1* with *SMARCA4* (P = 2.5×10^−4^) and *SMARCC2* (P = 7.1×10^−5^) in the G-401 cell line, all known members of SWI/SNF complexes and known to regulate transcriptional responses to hypoxia by modulating chromatin accessibility at HIF target genes.

**Figure 3:**
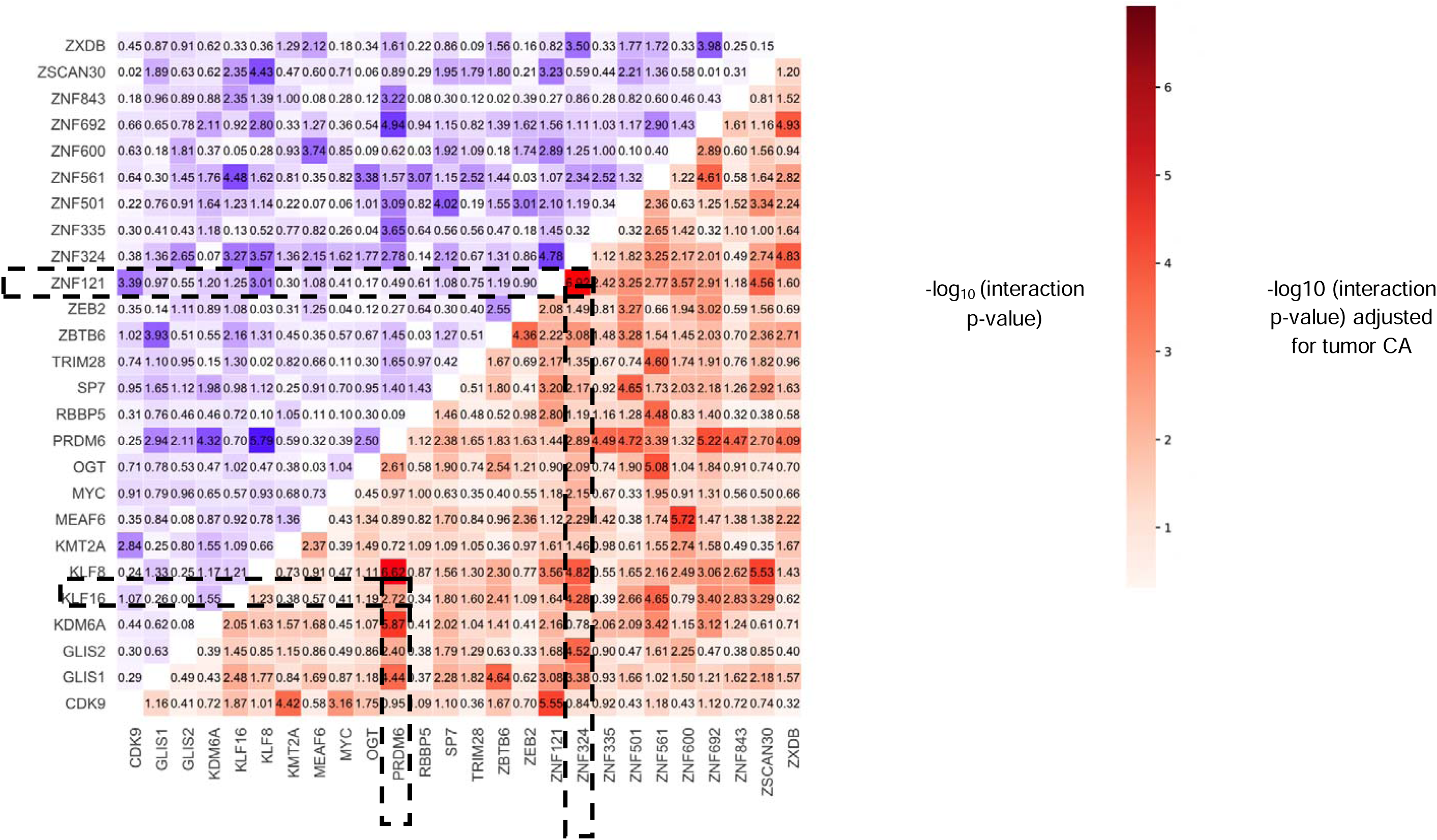
Co-occupancy of TFs. Lower right triangular plot showing the -log10(p-value) for co-occupancy using interaction model (See **Methods**) for 26 TFs which includes the pairs of TFs with strongest evidence of association with RCC. Two example associations of co-occupancy: PRDM6-KLF8 and ZNF121-ZNF324 are marked. Upper left triangular plot shows the -log10(p-value) for co-occupancy of the same pairs of TFs adjusted for tumor CA reported by Nassar et al.

### Cis-regulatory gene targets of risk TFs at RCC-associated loci

To delineate the *cis*-regulatory landscape and putative gene targets of risk transcription factors (TFs) in RCC, we queried enhancer-gene regulatory interactions across 17 distinct kidney-derived cell types and tissues using ENCODE-rE2G data^58^, which provides genome-wide enhancer-gene regulatory interaction maps generated using Activity-By-Contact (ABC)^59^ framework. We observed a total of 89 RCC-associated TF ChIP-seq profiles associated with at least one active chromatin mark (H3K4me2 and H3K27ac), linked to 235 unique target genes through enhancers containing 243 RCC-associated SNPs (P≤5×10⁻ ; **Supplementary Table 7**), encompassing 40 RCC associated regions. The number of enhancer-gene connections ranged from 9 to 181 per TF (mean= 85) with master regulators like *EP300* (181 targets), *SMARCA4* (161 targets) and *SMAD4* (159 targets) exhibiting higher number of unique *cis*-target genes, whereas other several key oncogenic regulators, such as *MYC* (119 targets), *EPAS1* (98 targets), *ARID2* (86 targets), and *ARNT* (28 targets), showed moderate levels of *cis*-target connectivity. A total of 145 genes had an ENCODE-rE2G connection in at least two cell types with RCC GWAS significant variants overlapping with RCC-related TFs. Of these, 54 were not previously identified in a previous transcriptome- or proteome-Wide association study (TWAS/PWAS)^60^. Among several RCC associated which have been previously reported on, we note that DPF3^10^ is the target of 55 different TFs across kidney cells including several members of SWI/SNF complex like PBRM1, SMARCA4, SMARCC2, BRD9 and SS18L1. In addition, we found 7 TFs linked to CCND1 (11q13.3)^30^, and 15 TFs linked to MYC (8q24.21)^8^.

### Trans-acting Effects of RCC Risk TFs on the Proteome

To investigate the downstream consequences of potential TF binding disruption at 106 RCC associated autosomal loci identified by Purdue et al^7^, we employed set-based regression approaches (See above and **Methods**) using variant-level summary statistics for 2,940 proteins reported by UKB-PPP^38^ on 34,557 individuals of European ancestry. We tested the association of the “TF-partitioned score” for each RCC-associated TF ChIP-seq profile, with 2,732 proteins that were at least 5Mb away from any of the RCC associated variants, representing *trans*-effects of potential TF binding disruption. Broadly this represents the association between effect of potential TF-binding altering variation with distal (*trans*) proteins. At a p-value threshold of 1.6×10^−7^, representing the Bonferroni correction threshold for 106 variants and 2,940 proteins, we identified 169 proteins *trans*-associated with the potential disruption of at least of one TF binding at RCC associated loci (**Figure 4**, **Supplementary Table 8**). Of these, 39 proteins were not individually associated with any single SNP highlighting the advantage of our approach in aggregating potentially weaker *trans*-effects, or the overall set of RCC-associated loci, highlighting the substantial heterogeneity within downstream effects of alterations in TF binding. The identified proteins include 9 tier-1 cancer genes^61^ and 39 candidate cancer driver genes overall^62^, significantly higher than expected by chance alone (P = 2.9×10^−4^). Of the identified proteins, 21 proteins were *trans*-associated to at least 10 TFs, indicating potential pleiotropic or converging effects of disruption in binding of multiple TFs and has high overlap with known drug targets (**Supplementary Figure 2**). This includes several potential cancer related genes like EDA2R (minimum p-value = 3.7×10^−31^), and multiple other members of the tumor necrosis factor receptor superfamily like TNFRSF11B (minimum p-value = 1.3×10^−12^) and TNFRSF8B (minimum p-value = 1.1×10^−13^). In contrast 25 proteins were *trans*-associated to one TF (**Table 1**), representing specific downstream targets. Among known key RCC and cancer genes, we identify KDR/VEGFR-2^63,64^ (minimum p-value = 4.8×10^−10^), target for anti-angiogenic therapies for RCC, to be trans-associated to RCC-loci at binding sites of 48 unique TFs, as well as AXL^65^ (minimum p-value = 2.9×10^−8^) and FBP1^66^ (minimum p-value = 3.9×10^−11^), associated to 8 and 6 unique TFs respectively. Among the TFs, zinc-finger TFs and master regulators (*EP300*, *TAF1* and others), had the highest number of downstream proteins, as expected. We also identified proteins that were consistently *trans*-associated with TFs of same family in multiple cell lines highlighting plausible downstream targets. For example, we identified 12 proteins *trans*-associated to at least one member of the HIF family, of which TLR3 (minimum p-value = 4.6×10^−17^) and ZP3 (minimum p-value = 1.0×10^−16^), both known prognostic markers for RCC^67–69^, were identified to be associated to *EPAS1* as well as *ARNT* in multiple cell lines. In fact, these two proteins were also trans-associated to RCC-related loci in *PBRM1* binding sites, highlighting the potential interaction between gene modules regulated by these TFs. The 169 identified proteins were enriched (**Supplementary Table 9**) in cytokine receptor interaction (P = 6.0×10^−10^), chemokine receptor binding (P = 1.4×10^−7^), PI3K-Akt signaling (P = 5.7×10^−4^) as well as targets of known TFs involved in RCC like c-JUN (P = 8.6×10^−6^) and c-FOS (P = 2.8×10^−3^).

**Figure 4:**
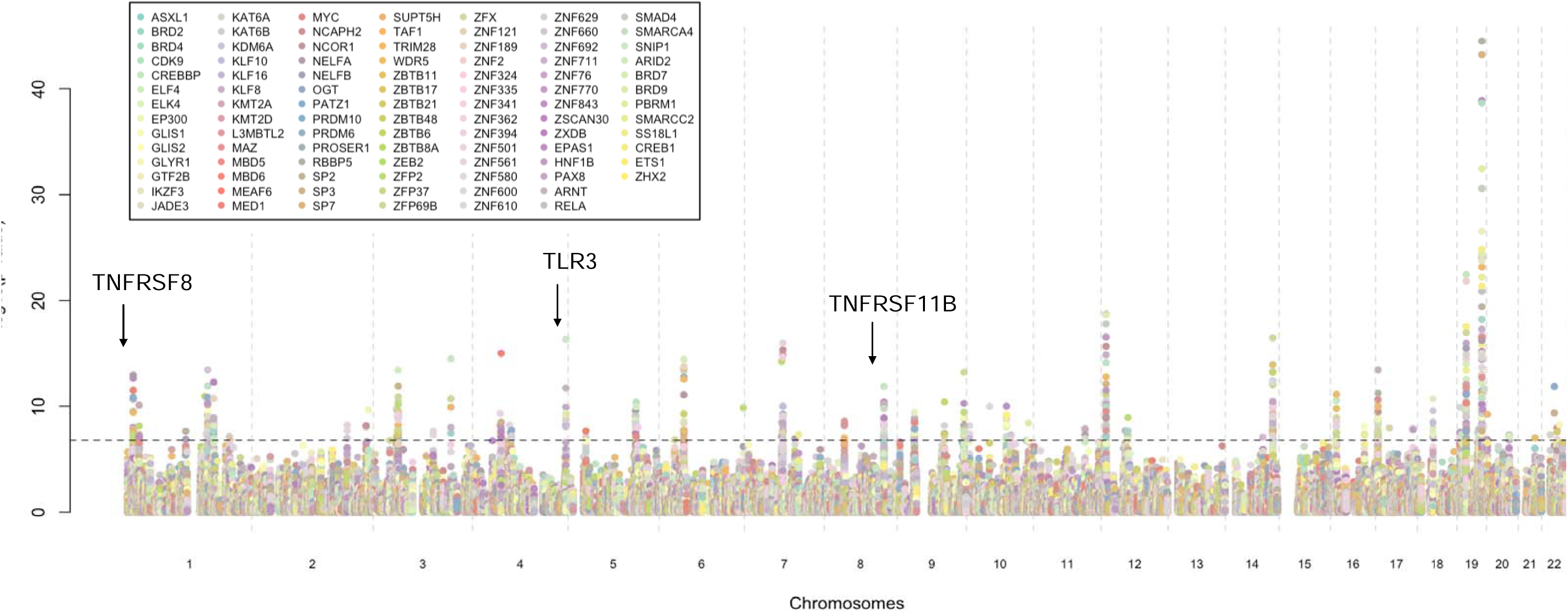
*trans*-effects of TF-binding disruption at RCC related loci. Manhattan plot showing the *trans*-association of 2,732 proteins profiled in UKB-PPP with each TF specific scores. Each dot represents -log10 p-value of association (vertical axis) for a protein (chromosome and transcription start site on horizontal axis), with a TF specific score and colored by the respective TF.

**Proteins trans** exactly 1 TF (See **Metho**

**Table 1: Proteins tran** exactly 1 TF (See Meth
| Protein | Transcription Factor | Cell Line | chromosome | Protein start | Protein end | p-value |
| --- | --- | --- | --- | --- | --- | --- |
| ADA2 | *ZFP37* | HEK293 | 22 | 17178790 | 17258235 | $4.74 \times 10^{-08}$ |
| ADAM15 | *ZNF362* | HEK293 | 1 | 155050566 | 155062775 | $1.16 \times 10^{-11}$ |
| AFP | *KLF10* | HEK293 | 4 | 73431138 | 73456174 | $5.73 \times 10^{-09}$ |
| ALPP | *EPAS1* | Renal cell carcinoma | 2 | 232378724 | 232382889 | $2.13 \times 10^{-10}$ |
| AXL | *ZFX* | HEK293 | 19 | 41219223 | 41261766 | $2.88 \times 10^{-08}$ |
| CD33 | *CREB1* | Kidney Cortex | 19 | 51225064 | 51243860 | $1.98 \times 10^{-10}$ |
| CEACAM16 | *ZSCAN30* | HEK293 | 19 | 44699151 | 44710718 | $6.99 \times 10^{-08}$ |
| COL2A1 | *ZNF580* | HEK293 | 12 | 47972967 | 48004554 | $2.00 \times 10^{-08}$ |
| CPXM2 | *WDR5* | HEK293 | 10 | 123706207 | 123940267 | $3.81 \times 10^{-09}$ |
| CXCL10 | *ZFP37* | HEK293 | 4 | 76021118 | 76023497 | $1.24 \times 10^{-07}$ |
| EPO | *EP300* | HEK293 | 7 | 100720468 | 100723700 | $1.10 \times 10^{-07}$ |
| FAS | *MBD5* | HEK293 | 10 | 88953813 | 89029605 | $4.76 \times 10^{-08}$ |
| FCRL5 | *ZFP37* | HEK293 | 1 | 157513377 | 157552515 | $4.02 \times 10^{-08}$ |
| ICAM5 | *EPAS1* | Renal cell carcinoma | 19 | 10289952 | 10296778 | $4.17 \times 10^{-08}$ |
| IDO1 | *ZFP37* | HEK293 | 8 | 39902275 | 39928790 | $2.55 \times 10^{-09}$ |
| IFNGR2 | *MEAF6* | HEK293 | 21 | 33403413 | 33479348 | $9.27 \times 10^{-09}$ |
| LAMB1 | *ZHX2* | Renal cell carcinoma | 7 | 107923799 | 108003213 | $4.66 \times 10^{-08}$ |
| LGALS9 | *ZFP37* | HEK293 | 17 | 27629798 | 27649560 | $1.09 \times 10^{-08}$ |
| LILRA3 | *CREB1* | Kidney Cortex | 19 | 54294163 | 54306461 | $2.24 \times 10^{-12}$ |
| MB | *SS18L1* | G-401 | 22 | 35606764 | 35637951 | $5.58 \times 10^{-09}$ |
| MSMB | *KMT2A* | HEK293 | 10 | 46033307 | 46048180 | $1.04 \times 10^{-10}$ |
| MZB1 | *ZFP37* | HEK293 | 5 | 139387467 | 139390081 | $9.46 \times 10^{-09}$ |
| NPC2 | *WDR5* | HEK293 | 14 | 74476192 | 74494177 | $8.04 \times 10^{-08}$ |
| NUDT2 | *ZBTB48* | HEK293 | 9 | 34329506 | 34343711 | $1.08 \times 10^{-07}$ |
| REN | *WDR5* | HEK293 | 1 | 204154819 | 204190324 | $6.82 \times 10^{-08}$ |
| SMOC2 | *WDR5* | HEK293 | 6 | 168441151 | 168673445 | $1.40 \times 10^{-10}$ |

## Discussion

This study represents to our knowledge the first large-scale agnostic scan of TF effects on RCC using publicly available GWAS summary statistics and ChIP-seq data. We identify 96 distinct TFs significantly associated with RCC risk which includes known regulators previously associated with RCC as well as master regulators, associated to multiple distinct cancers. In analyses restricted to RCC subtypes, we identified key regulators associated with ccRCC, which was qualitatively similar to RCC, while papRCC exhibited unique patterns, with TFs notably distinct from both RCC and ccRCC. Exploring TF co-occupancy, we uncover multiple cooperative interactions that are associated with RCC risk beyond the effect of individual TFs highlighting their complex interplay. Querying existing gene-enhancer connectivity maps at RCC associated loci, we identified *cis*-targets genes for each TF, potentially highlighting that certain TFs may function as specialized regulators with targeted specificity, while others may manifest broader regulatory functions with extensive target repertoires. We further characterize the downstream consequences of TF binding disruption at GWAS significant RCC loci by leveraging the largest available plasma proteomic data from the UK Biobank, identifying sets of proteins *trans*-regulated by altered TF binding, highlighting downstream targets for each TF.

Previous post-GWAS analyses with candidate TFs at selected GWAS loci identified specific mechanisms of action, notably implicating HIFs and SWI/SNF complex^8,10,30^. Our results provide an expanded list of potential TFs for mechanistic exploration, with additional candidate TFs at previously reported loci as well as on a genome-wide scale. While motif-level analysis is needed for further validation at specific loci, our results propose an initial list of candidate TFs that are associated with RCC susceptibility. Notably, the associations remained significant for majority of the TFs after adjusting for tumor CA, indicating cumulative effects of the variants in TF binding sites beyond CA. This includes several known and previously reported regulators of RCC pathogenesis like *ARNT* and *EPAS1*, essential components of the HIF family, *PAX8*, a lineage-specific TF, and *PBRM1*, a SWI/SNF complex subunit^25,27,44,46^. Further we also identify key master cancer regulators like *SMAD4*, *EP300*, *KAT6A* and *TAF1* which are associated with multiple cancers and are related to critical oncogenic processes^49,51,70^. By mapping TF binding to active chromatin regions, this study identifies regulatory hotspots consistently marked across RCC tumors, revealing how chromatin structure and TF activity converge to drive RCC pathogenesis. For several TFs, the overall effect of common variants present both at the binding sites of these TFs and active promoter/enhancer regions of the RCC tumor, have a marked stronger effect. For instance, *SMARCA4*, a core SWI/SNF complex component^71^, showed stronger associations at both promoter and enhancer regions, emphasizing its regulatory potential and its role in chromatin remodeling in RCC.

Our results highlight the critical role of TF co-occupancy and synergistic regulation in RCC, by identifying 220 significant interactions between TFs beyond their individual effects. For example, *PRDM6*, a transcriptional repressor linked to vascular differentiation, has significant interaction with *KLF8*, a zinc finger TF implicated in epithelial-to-mesenchymal transition (EMT) and invasion, potentially amplifying related pathways like promoting angiogenesis and metastasis, key features of RCC^72,73^. Similarly, the interaction between *BRD7* and *MYC* underscores the balance between tumor suppression and oncogenesis. *BRD7*, a SWI/SNF subunit, can act as a tumor suppressor. Its interaction with *MYC*, a potent oncogene, could suppress *MYC*-driven proliferation and metabolic reprogramming. Disruption of this interaction could enhance regulation of *MYC* activity, leading to aggressive tumor behavior^74^. A particularly striking finding is the interaction of *PBRM1* with SWI/SNF components *SMARCA4* and *SMARCC2*, emphasizing the role of chromatin remodeling in ccRCC. *PBRM1*, frequently mutated in RCC, modulates chromatin accessibility and HIF target gene expression^75^, crucial for angiogenesis, metabolism, and immune evasion. Dysregulation within the SWI/SNF complex impairs hypoxia responses, promoting tumor progression^76^. These underscore the importance of co-regulatory networks and chromatin remodeling in RCC but requires extensive laboratory investigation in future.

We further use a set-based regression approach to identify downstream *trans*-regulatory effects of TF binding alterations at RCC-associated loci, by integrating plasma proteomic data from UKB-PPP. Using TF-specific scores for GWAS significant RCC variants, our approach identifies *trans*-associated proteins, which can be interpreted as the molecular target which aggregates the effect of TF-binding alteration for RCC, highlighting it as a potential downstream target of the TF. We identify 169 proteins overall, including KDR/VEGFR-2, a central mediator of angiogenesis and a critical target of anti-angiogenic therapies^63,64^, reinforcing its pivotal role in RCC. Similarly, we identified proteins with known relevance to RCC like AXL, a target of cabozantinib, effective treatment for RCC^77,78^, and FBP1, a metabolic regulator frequently affected in clear cell RCC (ccRCC)^66^, highlighting how disrupted TF activity may drive downstream protein levels and oncogenic processes. We found synergistic roles of TF families, where effects of multiple TFs converge on the same proteins. For example, the *trans*-association of TLR3 and ZP3 with key TFs such as *EPAS1*, *ARNT* and *PBRM1* provides new insights into how genetic disruptions in hypoxia and chromatin regulation pathways converge on downstream protein targets. These proteins, both known prognostic markers for RCC, reflect the systemic impact of TF dysregulation on pathways governing inflammation, immune responses, and cellular stress adaptation in RCC^68,69^. Taken together the proteins were overrepresented in pathways important in cancer and specifically RCC, which highlights their potential for downstream targeting.

Our results underscore the central role of the SWI/SNF chromatin remodeling complex plays in RCC, through the regulation of chromatin accessibility and its interaction with HIF-regulated transcriptional programs. Notably a previous post-GWAS analysis identified components of this complex to be dysregulated by RCC-associated variants at 14q24.2^10^. Frequent mutations or dysregulation of core SWI/SNF components, are hallmarks of RCC, particularly in ccRCC^10,76^. Here we identify multiple members of the SWI/SNF complex to be associated to RCC like *PBRM1*, *SMARCC2* and *SMARCA4*. Notably *SMARCA4*, a core ATPase subunit of the SWI/SNF complex, exhibited stronger associations with RCC in active regulatory regions of RCC tumors, underscoring its role in enhancer activation. Moreover, the study identified interactions between SWI/SNF components such as *PBRM1*, *SMARCA4*, and *SMARCC2* and RCC-associated transcription factors, suggesting a synergistic regulatory network that integrates chromatin remodeling with transcriptional control of genes involved tumorigenesis. This complex plays a critical role in regulating transcriptional responses to hypoxia, a hallmark of ccRCC. In particular, loss of *PBRM1*, impairs the regulation of hypoxia-inducible factor (HIF) target genes, a key pathway activated in VHL-deficient tumors^75^. We identify proteins such as TLR3 and ZP3, *trans*-regulated by disruptions in binding of *PBRM1* as well as TFs of the HIF family. These can thus be thought of as potential molecular targets for SWI/SNF mediated HIF dysregulation associated to RCC.

There are several notable features in the statistical approach of the current study. This study conducts a scan of the TF landscape of RCC, by aggregating potentially weaker effects across the genome. By leveraging the largest RCC GWAS, our approach can improve statistical power to identify such weaker effects. Previous investigations have either been limited to analysis of candidate TFs or enrichment of binding at specific genomic regions. In contrast, our approach uses effects across the genome to identify the collective association of the variants at the binding sites for a TF.

The co-occupancy analysis builds on this approach, by identifying effects beyond those of individual TFs highlighting the role of cooperative regulation in the genetic architecture of RCC. Additionally, identifying the *trans*-regulatory consequences of TF binding disruption on the proteome can generate new insights into downstream consequences of TF binding alteration. Our approach aggregates the effects multiple variants which can potentially alter the binding of a given TF and identify the proteins *trans*-associated with this group of variants. Not only this approach can improve statistical power by cumulating the effects of multiple variants but also offers unique interpretation of the protein as the potential target of the alteration in binding of a corresponding TF, highlighting the distinct mechanisms regulated by specific subsets of RCC related variants.

Our study has limitations which should be considered in interpreting these results. Firstly, most of the TF ChIP-seq data used in this study come from non-RCC kidney cell lines (e.g., HEK293), which may not fully capture the distinctive regulatory architecture characteristic of RCC. Thus, these associations need to be reevaluated in cancer cell lines for further validation. Second, the observed difference between results for ccRCC and papRCC can be driven potentially by the large difference in number of cases (16,321 for ccRCC and 2,193 for papRCC) for each of them. Third, we have reported several TF co-occupancy interactions, but the biological mechanisms behind these interactions remain unclear. While querying established interaction databases reveals evidence of direct TF-TF interactions, alternative mechanisms may also contribute to such patterns. Additional analyses incorporating Hi-C data and motif analysis is needed to provide more insights into whether these interactions reflect cooperative regulation within topologically associating domains and enhancer-promoter contact sites or represent or independent binding events at separate variants within the same loci.

In conclusion, this study demonstrates the value of integrative multi-omic analysis to uncover the elements of the altered regulatory mechanisms underlying RCC and its subtypes. By dissecting the transcriptional regulatory landscape of RCC, we have identified key TFs, cooperative interactions, and downstream pathways that mediate genetic risk and disease pathogenesis. However, the majority of these findings are based on broad patterns of association. As a next step it is necessary to refine such results through further in silico analysis using larger epigenomic and transcriptomic data from RCC tumors, as well as functional screening approaches such as CRISPR-based perturbation assays as well detailed experimental validation and laboratory investigation. These findings provide a foundation for further exploration and validation, with the potential to transform our understanding of RCC biology and inform the development of new diagnostic, prognostic, and therapeutic strategies.

## Data Availability

This analysis used publicly available GWAS summary statistics. GWAS summary statistics for RCC, ccRCC and papRCC are publicly available as outlined in Purdue et al. on dbGaP (<u>phs003505.v1.p1</u>) and also in the GWAS Catalog (<u>GCST90320043 – GCST90320065</u>). TF ChIP-seq experiments were curated from ENCODE: https://www.encodeproject.org/ and ChIP-Atlas: https://chip-atlas.org/. Epigenomic profiling of RCC tumors generated by Nassar et al^53^ were downloaded from Gene Expression Omnibus (GEO) accession number <u>GSE188486</u>. ChromHMM: https://egg2.wustl.edu/roadmap/data/ (E086 cell). UKB-PPP: https://www.synapse.org/Synapse:syn51364943/wiki/622119. Gene-set enrichment was conducted using gProfilerR: https://biit.cs.ut.ee/gprofiler/gost. Drug targets analysis was performed with GSCA: https://guolab.wchscu.cn/GSCA/. List of cancer genes were curated from COSMIC: https://cancer.sanger.ac.uk/cosmic and NCG: http://network-cancer-genes.org/. ENCODE-rE2G enhancer to gene connectivity maps were downloaded from publicly available datasets (https://www.encodeproject.org/search/?searchTerm=ENCODE-rE2G). ATAC-seq from TCGA-KIRC: https://gdc.cancer.gov/about-data/publications/ATACseq-AWG.

## Code Availability

Custom code used for data analysis are posted on GitHub: https://github.com/chhetribsurya/RCC-TF-analysis.

## Acknowledgements.

SB and AB were supported by National Institutes of Health Grant R35GM139580. The remaining authors were supported by the Intramural Research Program of the National Cancer Institute, National Institutes of Health.

## Methods

### Genome-Wide Association Study (GWAS) of RCC and its subtypes

We obtained summary statistics from the current largest multi-ancestry RCC GWAS meta-analysis conducted by Purdue et al^7^, which included 29,020 cases and 835,670 controls for overall RCC, 16,231 cases and 743,479 controls for ccRCC and 2,193 cases and 740,816 controls for papRCC. The details of the preprocessing and GWAS analyses have been described previously. For each SNP, we obtained the z-value (probit transformed p-value; See below), quantifying its association with the outcome (RCC or its subtypes) in gaussian scale. The authors also report 106 independent autosomal loci associated to RCC, the corresponding index SNPs and effect sizes (log odds ratio), which were used to identify trans-effects of TF binding disruption at these loci, across the proteome (See below).

### Genome Partitioning

We partitioned the genome into approximately independent LD blocks. Each LD block was defined as a 2Mb genomic region centered on the sentinel GWAS variant, which was identified as the SNP with the lowest p-value in each risk locus. This partitioning scheme ensures that variants within each LD block are in strong LD with the sentinel variant and that blocks are approximately independent from each other.

#### Transcription Factor Binding Profiles

Genome-wide binding profiles for 449 TFs were obtained from publicly available ChIP-seq datasets, including ENCODE^35^, ChIP-atlas^36^ and various studies deposited in the Gene Expression Omnibus (GEO) for relevant cell types, such as Human embryonic kidney, renal proximal tubule epithelial cells and RCC cell lines (e.g., HEK293, 786-O, TTC1240, G-401). ChIP-seq peak files were downloaded and processed peak calls were converted to hg38 coordinates using the UCSC LiftOver tool. For each TF, we then defined “occupancy” for every SNP in the GWAS summary statistics, as a binary variable taking values 1 when a SNP is within a 100Kb window around a ChIP-seq peak and 0 otherwise.

#### Chromatin Accessibility

We curated RCC tumor chromatin accessibility (CA) data from two sources: (1) ATAC-seq data from the cancer genome atlas (TCGA) kidney clear cell carcinoma (KIRC) across 16 tumors^51^ and (2) ATAC-seq data generated by Nassar et al^52^ in six RCC tumors including clear cell, chromophobe and papillary tumors across different stages and grades.

#### Epigenomic Profiling

We used histone modification (H3K27ac and H3K4me2) profiling from Nassar et al^52^ across 42 patients with different subtypes of RCC. For each histone subtype and histone modification, we defined (1) consensus peaks: denoting the regions which were histone modification ChIP-seq peaks across all the patients with the RCC subtype and (2) union peaks: denoting the regions which were histone modification ChIP-seq peaks in at least one the patients with the RCC subtype. Thus, we defined 4 sets of annotations: consensus and union peaks for ccRCC and papRCC subtypes and H3K4me2 and H3K27ac histone marks.

These were used to integrate with TF ChIP-seq peaks to identify regions which were in TF ChIP-seq peaks as well as histone marks in a particular RCC subtype.

#### Protein Expression Data

UK Biobank Pharma Proteomics Project (UKB-PPP) used the Olink platform to measure 2,940 protein analytes in 54,306 individuals across 4 ancestry groups and have made the summary statistics publicly available. The normalization and preprocessing pipeline have been reported previously. We used the summary statistics from the discovery cohort of about 34,557 individuals of European ancestry here, to identify effects of TF occupancy disruption at RCC associated loci, across the proteome. In particular, to identify trans-effects we used data on 2,732 proteins which were at least 5Mb away from all of the 106 RCC related index variants reported by Purdue et al.

### Statistical Model

*<u>Association with TF occupancy.</u>* The primary intuition of our approach is to identify association between the occupancy of TFs and z-values (See above) of SNPs. If there is a significant association, we can declare the corresponding TF to be associated with risk of the disease overall. To test for association between RCC GWAS signals and TF binding sites, we employed a linear mixed model:

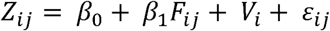

where Z_ij_ is the Z-score for the j^th^ SNP in the i^th^ LD block, calculated from the GWAS p-values of each SNPs as 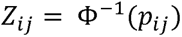, where *p_ij_* is *F_ij_* is the “occupancy” of the TF of interest, a variable taking values 1 when the j^th^ SNP in the i^th^ LD block overlaps with a ChIP-seq peak for the TF of interest and 0 otherwise; *v_i_* is a random effect term for the i^th^ LD block, capturing block-specific correlation and *ε_ij_* is the residual error term. The fixed effect coefficient *β*_1_ represents the association of TF occupancy with RCC risk, with a positive value indicating that SNPs overlapping TF peaks have larger Z-scores than expected by chance. We test whether *H_0_*:*β*_1_=0 against *H_0_*:*β*_1_>0 and the resulting p-value indicates the strength of association between the TF and RCC risk. We use a p-value threshold of 1×10^−04^ corresponding to Bonferroni correction for 449 TF ChIP-seq profiles included in the analysis. It is to be noted that such linear mixed model-based approach has been used in the past by Wen et al for breast cancer. However, our current approach using z-values derived from probit transformation would produce better calibration of false positives theoretically.

#### Comparison with other methods

We compared our results with that from stratified LD score regression (S-LDSC) and bootstrap enrichment. S-LDSC partitions the genome into functional annotations (e.g., TF binding sites according to occupancy of SNPs described above) and estimates the contribution of each annotation to disease heritability while accounting for LD structure. We computed LD scores for each TF binding annotation using the 1000 Genomes EUR reference panel and applied S-LDSC to the RCC GWAS summary statistics. Enrichment was assessed using the proportion of heritability explained by each TF annotation divided by the proportion of SNPs in the annotation, after accounting for the baseline annotations. For bootstrap enrichment, we investigated the excess overlap between 106 autosomal GWAS significant RCC sentinel SNPs and a TF ChIP-seq profile of interest. First, we identified all SNPs in strong LD (*r*^2^ ≥ 0.8) with the index SNP at each of the 106 RCC-associated locus using LD extracted from 1000 Genomes EUR reference panel. We defined overlap as a locus coinciding with peaks in two or more of the five ChIP-seq datasets of HIF-binding sites. We defined overlap as a locus coinciding with TF ChIP-seq peaks and performed a bootstrapping analysis by randomly shuffling the GWAS significant loci 500,000 times across the genome and at least 5 Mb away from the originally identified loci, evaluating statistical significance against the null hypothesis that the RCC susceptibility loci are not enriched for TF ChIP-seq peaks.

#### Adjusting for tumor chromatin accessibility (CA)

Association of a TF ChIP-seq profile with RCC (or its subtypes) can, in part, be driven by the variation in chromatin accessibility (CA) along the genome. To test for association between RCC GWAS signals and TF ChIP-seq profiles, controlling for tumor CA, we modified the model as:

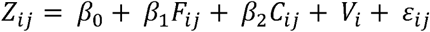

where C_ij_ is a variable taking values 1 when the j^th^ SNP in the i^th^ LD block lies within a CA region. As KIRC and (2) that reported by Nasser et al. Under this model, β_1_ reflects the effect of the TF ChIP-seq profile mentioned above, we used two sources to define CA regions in tumor using ATAC-seq data from (1) TCGA-on RCC beyond the effect of tumor CA. For the TF ChIP-seq profiles identified to be significant in previous analysis, we apply this model to evaluate the whether the associations are significant after adjusting for tumor CA.

#### Integrating epigenomic profiling with TF occupancy

To test for association between RCC GWAS signals and TF binding sites, restricted to a genomic annotation derived from epigenomic profiling (See above), we employed a similar linear mixed model:

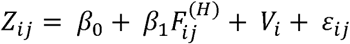

where 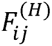 is the occupancy of the TF of interest, a variable taking values 1 when the j^th^ SNP in the i^th^ LD block overlaps with both the ChIP-seq peak for the TF of interest and genomic annotation of interest, and 0 otherwise. For genomic annotation we used consensus and union peaks for H3K4me2 and H3K27ac in each overall RCC, while peaks for papRCC were used to analyze summary statistics for papRCC. Here β^1^ subtype ccRCC and papRCC. Peaks for ccRCC was used to analyze summary statistics for ccRCC as well as represents the association of TF occupancy with RCC risk, restricted to the genomic annotation, with a significant positive value indicating that SNPs overlapping TF peaks and genomic annotation, have larger Z-scores than expected by chance.

#### Association with TF co-occupancy

To test for association between RCC GWAS signals and co-occupancy for two TFs, we incorporated an interaction effect as follows:

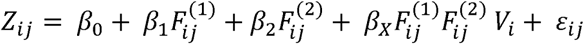

where 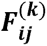 is the occupancy of the j^th^ SNP in the i^th^ LD block for the k^th^ TF, β_k_ is the independent effect of the k^th^ TF and β_X_ is the interaction effect. We test whether ***H_0_*:*β*_x_>0** and if significant, it can be interpreted as a significant effect of co-occupancy of the TFs on RCC, beyond the effect of individual TFs. We evaluate the robustness of the findings, by adjusting the model for tumor CA, as an additional covariate as above, and report which associations remain significant after the adjustment.

#### Trans-effects of TFs at RCC-associated loci on the proteome

To identify the trans-effects of TF binding disruption at RCC-associated GWAS loci, we integrated plasma proteomic data from the UK Biobank Pharma Proteomics Project (UKB-PPP) with the 106 independent loci associated to RCC identified by Purdue et al. For each TF, we defined a "TF-specific subset" of variants among the 106 index SNPs, by using the following criteria:

- The index SNP overlaps with a ChIP-seq peak, or,
- Any SNP with LD (*r*^2^ ≥ 0.75) with the index SNP overlaps with the ChIP-seq peak.

Representing the (imputed) genotypes of the i^th^ SNP in "TF-specific subset" as G_sub:i_, we then define TF-partitioned score as,F_sub:i=_β_i_G_sub:ii_ a vector of length *n* where β_i_ is the estimated effect size (log odds ratio) of the corresponding SNP on overall RCC, as reported by Purdue et al *n* and is the sample size for the proteomic data.p_ki_, a vector values represents the contribution of the i^th^ SNP to RCC risk for each individual. For a given protein, the association between normalized plasma protein levels p_k_ and these TF-partitioned scores across *n* individuals, using a set-based regression approach:

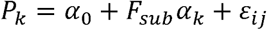

where *F*_sub_ is the matrix of TF-partitioned scores for all the TF-specific variants (columns) across *n* individuals (rows). α_k_ is a random effect following *N(0,τ*^2^*Ψ)*, with τ = 0 under no association between the TF and TF-specific scores and *Ψ= 0* denoting the relation (kernel matrix) between the effects of the TF specific scores on the protein. A significantly non-zero α_k_ indicates a significant effect of TF-specific subsets on protein levels, highlighting potential downstream targets of TF binding disruption.

We tested associations using variance component tests^39–41^, under two assumptions: (1) uniform effects: the effect of each TF-partitioned score on the protein level is identical (equivalent to Ψ = 11’ or all elements of α_k_ being same) and (2) independent effects: the effect of each TF-partitioned score on the protein level is independently distributed (equivalent to Ψ= I, or all elements of α_k_ are independent).

(1) Under uniform effects (*Ψ =11’*) assumption, we test the association between TF-partitioned scores and protein level using the test statistic:

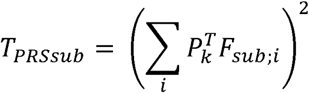

which follows a mixture of chi-squares under the null hypothesis of no association. In particular, this amounts to testing whether the sub-PRS corresponding to the variants in TF-specific subset has significant association with the protein. Broadly this is akin to performing “burden test” using TF-specific scores, thus amounting to testing the association between the cumulative effect of TF binding disruption and the protein level. We thus refer to this as PRS_sub_.

(2) Under independent effects (*Ψ= I*) assumption we test the association between TF-partitioned scores and protein level using the test statistic.

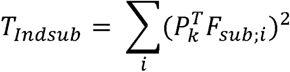

Similarly, this is same as performing “sequence kernel association test (SKAT)” using TF-specific scores, indicating independent effects of the TF specific variants on the protein level. We thus refer to this as Ind_sub_. We expect Indsub to have higher number of discoveries than PRSsub due to the nature of the assumptions.

Of note, both of these association test can be computed using variant level summary statistics for the proteins as reported by UKB-PPP using the simple algebraic relation:

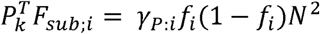

where γ_P:i_ is the effect of ith SNP on the protein P, extracted from the GWAS of each protein reported by UKB-PPP, *f*_i_ is its minor allele frequency and *N* is the total sample size on which the GWAS of protein P was conducted, which is 34,557 for UKB-PPP. Corresponding to each TF, we extract the proteins that have distal (> 5Mb) transcription start sites from each of the SNPs constituting its TF-specific variants. We test the association of all such distal proteins with the corresponding TF-partitioned score using both PRS_sub_ and Ind_sub_. Any association with p-value < 1.6×10^−7^ is considered to be a significant trans-effect of the TF binding alteration on the protein. Intuitively, this can be viewed as convergent effect of potential TF binding alterations at RCC related SNPs on a protein and thus can be interpreted as a downstream target for the corresponding TF broadly.

### Post-hoc analysis

#### Genomic Annotations

For each TF ChIP-seq identified to be significantly associated with RCC or its subtypes (See above), we annotated the variants in the ChIP-seq peaks using epigenetic data from Roadmap projects. For each genetic variant, we characterized whether variants mapped to functional regions (i.e., promoter or enhancer) using ChromHMM 18-state model annotation in fetal kidney (ROADMAP E086; See **Data availability**).

#### Gene-set enrichment

We performed standard pathway/gene-set overrepresentation analysis using gProfiler software (See **Data availability**) to test whether a given set of genes/proteins were enriched in predefined set of genes. This analysis was done using (1) the unique TFs identified to be significantly associated to RCC and (2) proteins identified to be associated to at least one TF-specific score.

#### Allelic imbalance

To assess the allele-specific binding patterns of RCC-TFs, we implemented an allelic imbalance analysis pipeline for 7 randomly selected TF ChIP-seq profiles from HEK293 cell line that were significantly associated with RCC. ChIP-seq alignment files in BAM format were obtained from the ENCODE portal (See **Data availability**) for RCC-TFs of interest. These BAM files, containing aligned sequencing reads, were processed using the python-based pysam tools (version 0.23.0). We processed SNP genomic coordinates (with effect and non-effect allele information) associated with RCC to quantify allele-specific read counts. For read counting, pysam’s pileup function algorithm was employed to generate base-level resolution coverage information at each SNP position, systematically examining the genomic region surrounding each SNP. Default base quality and mapping quality filters were applied to ensure accurate read assignments. The reads supporting each allele at RCC-associated SNP loci was counted, generating a comprehensive allele count matrix, and additionally, a minimum read depth threshold of ≥ 5 reads per SNP was applied to ensure sufficient sequencing coverage. For the statistical evaluation of allelic imbalance, we employed a standard binomial test using scipy stats under the null hypothesis of equal allelic distribution (P = 0.5), assuming no preference for either allele. A two-sided alternative hypothesis was used to detect significant deviations from the expected 50:50 allelic ratio by comparing A1 (effect allele) versus A2 (non-effect allele). A binomial test is performed to assess deviation from expected allelic ratios, and a p-value is computed for each SNP to determine whether one allele is overrepresented. This approach enables the identification of TFs that show allele-specific occupancy or preferential binding to a specific allele, providing insights into the potential regulatory impact of genetic variants on RCC-TF binding.

#### Protein interactions

Protein interaction data was curated from 3 separate sources including STRING Db, CORUM and BioGRID to evaluate whether identified interactions between TF co-occupancy pairs reflect physical interactions.

