## Supplementary Figure for "Integrative multi-omic analysis identifies key transcription factors and target proteins in renal cell carcinoma and its subtypes"

**Supplementary Figures**

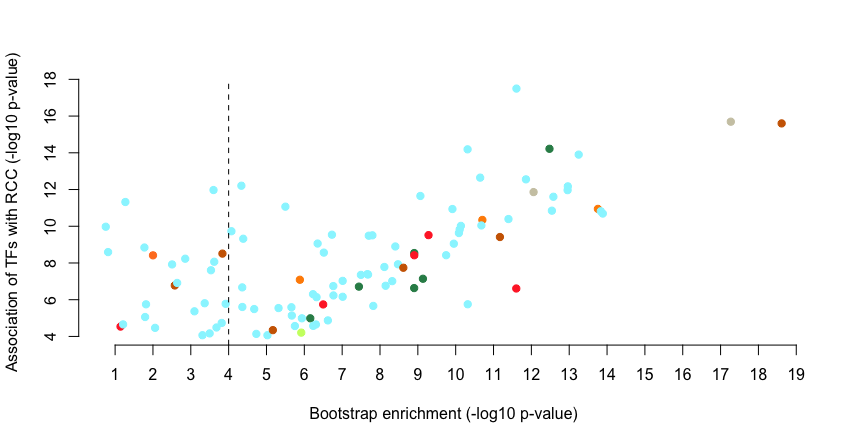

**LMM vs Bootstrap**

**LMM vs S-LDSC**

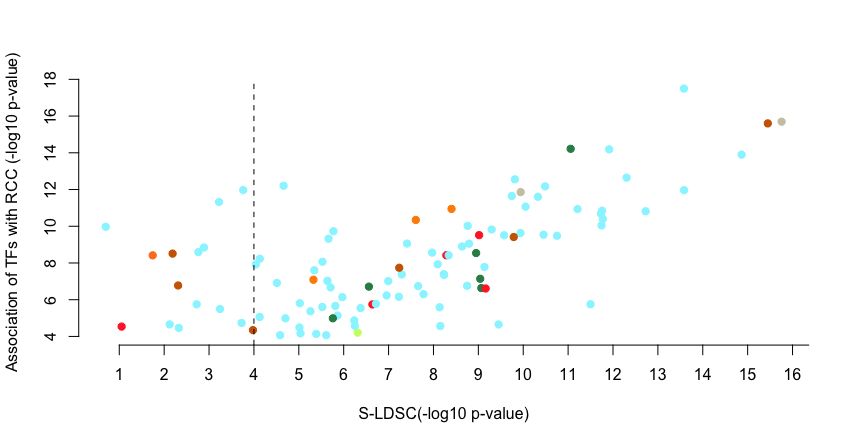

**Figure S1:** Comparison of association of TFs with RCC (colored by cell lines) using the mixed model-based approach (See **Methods**) against that with Bootstrap enrichment (upper panel) and stratified LD score regression (S-LDSC; lower panel).

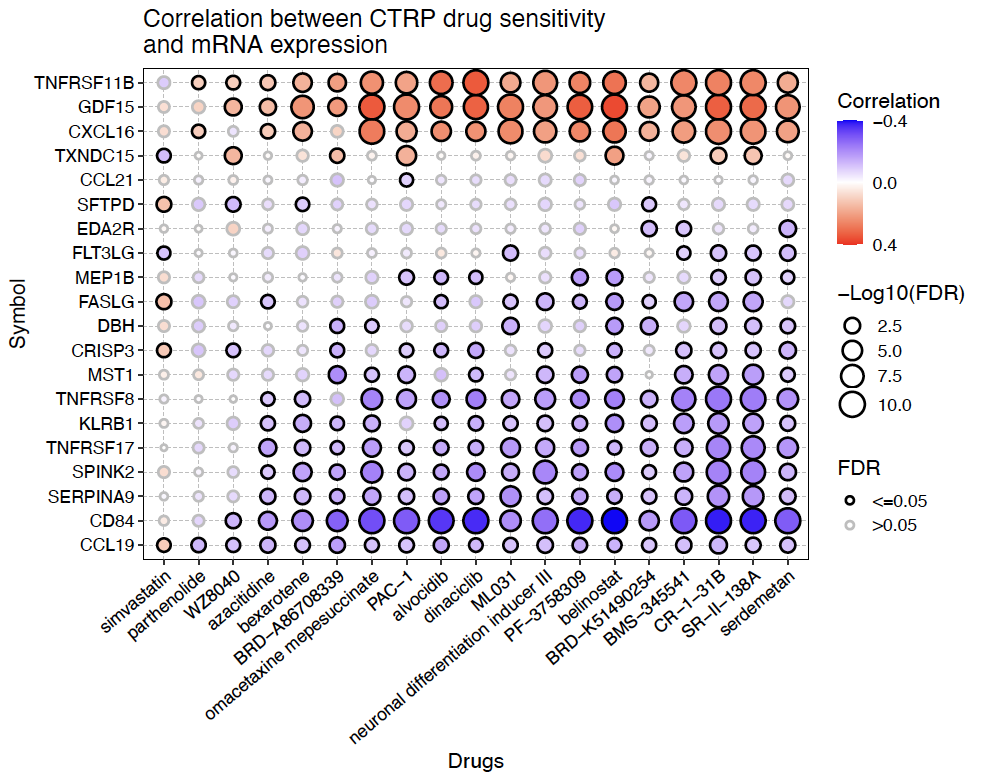

**Figure S2:** Bubble plot showing the correlation between gene expressions corresponding to 21 proteins trans-associated at least 10 TFs and IC50 for different drugs. P-values for the test of correlation adjusted for false discovery rate. Data and plot generated from Gene-set Cancer Analysis web portal(GSCA: [www.guolab.wchscu.cn/GSCA](http://www.guolab.wchscu.cn/GSCA))
